# Study on the expression of RAD51 in non-small cell lung cancer based on bioinformatics

**DOI:** 10.1101/2023.01.29.526152

**Authors:** Jinghong Wu, XianYu Zhang, Yanmei Zhang, Fan Gao, Guangyan Wang

**Author notes:** These authors contributed equally to this work (Jinghong Wu, XianYu Zhang).

## Abstract

**Objective:** RAD51 is a DNA repair protein, which participates in the resistance of tumor cells to radiotherapy/chemotherapy and reduces the therapeutic effect. Based on the research status of RAD51 at home and abroad and the analysis of online databases, the purpose of this study was to explore the relationship between RAD51 expression and clinical patient survival and prognosis. It is expected to provide a new theoretical basis for the clinical treatment of lung cancer patients, help identify new molecular markers, and provide new targets for the biological therapy of lung cancer patients.

**Methods:** the RNA Seq data of NSCLC in TCGA database were downloaded, and the expression of RAD51 gene in NSCLC and normal tissues were analyzed by R studio software. Clinical correlation analysis revealed its correlation with the clinicopathological characteristics of non-small cell lung cancer. Survival analysis was used to evaluate the relationship between the expression level and the prognosis of patients. CIBERSORT and TIMER were used to evaluate the correlation between the expression level of CIBERSORT and immune cells in the tumor microenvironment. The protein expression level of RAD51 in non-small cell lung cancer was evaluated by HPA.

**Results:** RAD51 was highly expressed in lung cancer (p<0.05), which was significantly associated with poor prognosis of lung adenocarcinoma patients (p=0.0026), but not with lung squamous cell carcinoma (p=0.76). The expression level of RAD51 mRNA was associated with different pathological stages of lung adenocarcinoma (p=0.000528), but not with different pathological stages of lung squamous cell carcinoma (p=0.326). RAD51 was positively correlated with the expression of TP53, BRAF, EGFR, MYC, PD-L1, and KRAS (p<0.001). In lung adenocarcinoma, lung cancer cells were positively correlated with CD4+memory T cells, CD8+T cells, and M1 macrophages (p<0.001). In lung squamous cell carcinoma, tumor cells were positively correlated with M1 macrophages (p<0.05), but not with CD4+T memory cells, CD8+T cells, M2 macrophages, and Tregs cells (p>0.05). The HPA database indicated that RAD51 protein was positive in non-small cell lung cancer.

**Conclusion:** RAD51 is highly expressed in non-small cell lung cancer and is associated with poor prognosis. RAD51 can be used as a biomarker related to the prognosis of non-small cell lung cancer and is expected to become a target for the diagnosis and treatment of non-small cell lung cancer.

---

Lung cancer remains the most common cause of cancer and cancer death worldwide [1]. More than 50% of patients die within the first year of lung cancer diagnosis, and the 5-year survival rate is less than 20%[2]. According to the latest survey, there were 787,000 new cases of lung cancer in China, with an incidence rate of 57.26/100,000 and a death rate of 63,100, respectively, with a mortality rate of 45.87/100,000. Compared with 2010, the number of new cases increased by 25.56%[3]. So far, smoking is the main risk factor for lung cancer, followed by pollution, occupational carcinogens, genetic susceptibility, lifestyle habits, and alcohol consumption [2, 4]. lung cancer is mainly divided into small cell lung cancer (SCLC) and nonsmall cell lung cancer (NSCLC), among which 85% of lung cancer cases are NSCLC, adenocarcinoma, LUAD) and lung squamous cell carcinoma (LUSC) are the most common subtypes [2]. According to the latest classification of lung cancer by the World Health Organization (WHO), lung cancer is mainly divided into three types: adenocarcinoma, squamous cell carcinoma, and neuroendocrine tumor. In addition, lung adenocarcinomas are classified into low-grade (squamous), intermediate (acinar and papillary), and high-grade (solid and micropapillary) types based on prognosis and excised specimens [5]. Depending on the pathological type and clinical stage of lung cancer, radiotherapy, chemotherapy, or surgical resection can be selected. Although great progress has been made in the treatment of lung cancer, the survival rate of patients is still low and radiotherapy/chemotherapy tolerance occurs in the late stage of treatment. Therefore, it is necessary to develop new therapeutic targets to improve the cure rate of lung cancer.

RAD51 is a DNA repair protein [6], which can realize accurate and timely DNA repair. The homologous recombination (HR) differs from RAD51 and is a major mechanism of DNA double-strand repair and a contributor to the induction of tumor tolerance to radiotherapy and chemotherapy. Vp16-induced DNA double-strand break repair mediates etoposide tolerance in NSCLC through RAD51-dependent homologous recombination and DNA-PKCs-dependent non-homologous recombination repair [7], and RAD51 induces chemotherapy tolerance in breast cancer and chronic lymphocytic leukemia by activating the HR pathway [8, 9]. RAD51 promotes aerobic glycolysis by up-regulating HIF1a protein to promote the development of pancreatic cancer. RAD51 inhibitors can enhance the blocking of cellular glycolysis and increase the therapeutic effect on leukemia [10, 11]. In addition, RAD51 has been shown to promote epithelial-mesenchymal transition in esophageal squamous cell carcinoma and prostate cancer via p38/Akt/Snail and EGFR-Erk1/2/Akt signaling pathways. EMT) and distal metastasis [12, 13]. RAD51 mediates tumor metastasis through a variety of mechanisms, which will become a new therapeutic target to inhibit tumor progression and become one of the key targets for the treatment of NSCLC.

This study analyzed the relationship between transcription level, protein expression, and prognosis of RAD51 in NSCLC from a large sample level through TCGA and related databases, to provide clues and ideas for the mechanism of action of RAD51 in the occurrence and development of NSCLC, and provide theoretical basis and scientific basis for the clinical treatment of radiotherapy/chemotherapy resistance in non-small cell lung cancer.

## 1. Materials and methods

### 1.1 Data Collection

Joint TCGA database (https://portal.gdc.cancer.gov/) and analysis RAD51 GEPIA database (http://gepia2.cancer-pku.cn/), expressed in a variety of tumors in the at the same time for lung adenocarcinoma and clinical data of lung squamous carcinoma.

### 1.2 The expression difference of RAD51 in NSCLC and paracancer tissues in TCGA database was analyzed

Non-parametric rank Test (Wilcox Test) and R studio software Beeswarm package were used to analyze the differences in expression of RAD51 mRNA obtained from the TCGA database in NSCLC and para-cancer tissues. p< 0.05 was considered statistically significant.

### 1.3 Analysis of the relationship between RAD51 expression in TCGA database and prognosis of NSCLC patients

According to NSCLC typing, RAD51mRNA expression levels of lung adenocarcinoma and lung squamous cell carcinoma were obtained from the TCGA database, and survival analysis was performed using Kaplan-Meier curve by R packet survival and survminer. p< 0.05 was considered statistically significant.

### 1.4 Correlation analysis between RAD51 and TP53, KRAS, EGFR, BRAF, MYC and ALK

The correlation between RAD51 and the above genes was analyzed by using the GEPIA database, and Pearson was selected for analysis. The results were presented as scatter plots. p< 0.05 was considered statistically significant.

### 1.5 Relationship between RAD51 and tumor-infiltrating immune cells in NSCLC tumor microenvironment (TME)

Through CIBERSORT (http://cibersort.stanford.edu/) joint RAD51 TIMER database analysis and infiltration of immune cells in the TME expression differences in NSCLC, the results displayed in a scatter plot. p< 0.05 was considered statistically significant.

### 1.6 Protein expression difference of RAD51 in human normal lung tissue and NSCLC

The expression differences of RAD51 gene in human normal lung tissue, lung adenocarcinoma and lung squamous cell carcinoma were obtained from HPA database (www.proteinatlas.org).

### 1.7 Statistical Methods

Wilcox Test was used to detect the difference of RAD51 gene expression between liver cancer and adjacent tissues. The relationship between RAD51 gene expression and prognosis of patients with non-small cell lung cancer was expressed by Kaplan-Meier curve. The clinical data were analyzed by multivariate Logistic regression. The correlation between RAD51 and gene was analyzed by Pearson. p< 0.05 was considered statistically significant.

## 2 Results

### 2.1 Expression difference and clinical correlation analysis of RAD51 in various cancers and adjacent tissues in GEPIA database and TCGA database

In the GEPIA database, RAD51 expression was elevated in a variety of tumors except acute myeloid leukemia. By analyzing the TCGA database, it was found that the expression of RAD51 in lung adenocarcinoma and lung squamous cell carcinoma was higher than that in the adjacent tissues, the difference was statistically significant (p< 0.05). The analysis of clinical data showed that RAD51 was not correlated with age, sex or race of patients with lung adenocarcinoma (p> 0.05), RAD51 was correlated with gender in lung squamous cell carcinoma (p< 0.05) (Figure 1, Table 1-2).

**Table 1.**
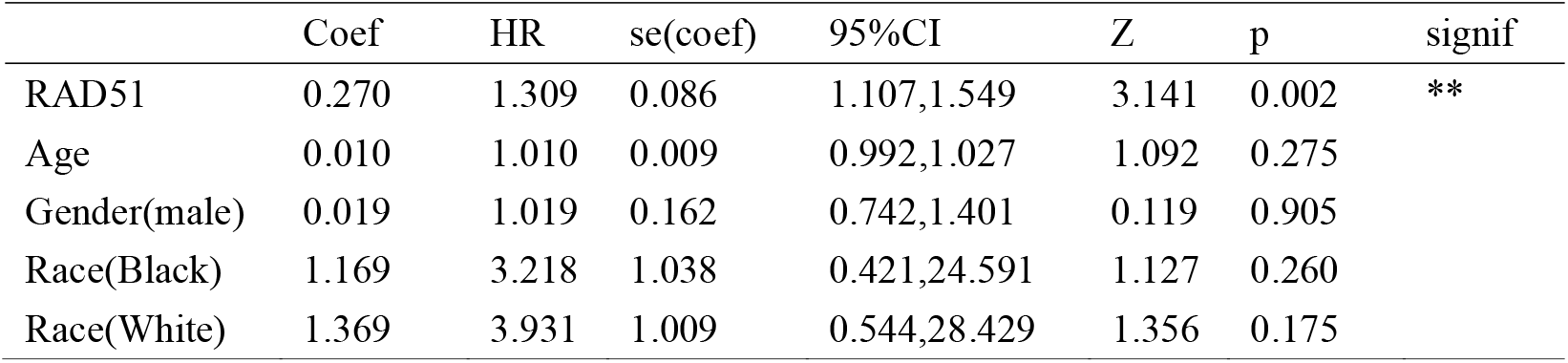
Multivariate regression analysis of lung adenocarcinoma

**Table 2.** Multivariate regression analysis of lung squamous cell carcinoma

|  | Coef | HR | se(coef) | 95% CI | Z | p | signif |
| --- | --- | --- | --- | --- | --- | --- | --- |
| RAD51 | -0.251 | 0.778 | 0.118 | 0.617,0.982 | -2.116 | 0.034 | * |
| Age | 0.014 | 1.014 | 0.009 | 0.996,1.033 | 1.535 | 0.125 |  |
| Gender(male) | 0.528 | 1.695 | 1.194 | 1.160,2.478 | 2.726 | 0.006 | ** |
| Race(Black) | 0.091 | 1.096 | 0.611 | 0.331,3.360 | 0.149 | 0.881 |  |
| Race(White) | -0.397 | 0.672 | 0.566 | 0.222,2.037 | -0.702 | 0.483 |  |

**Fig. 1.**
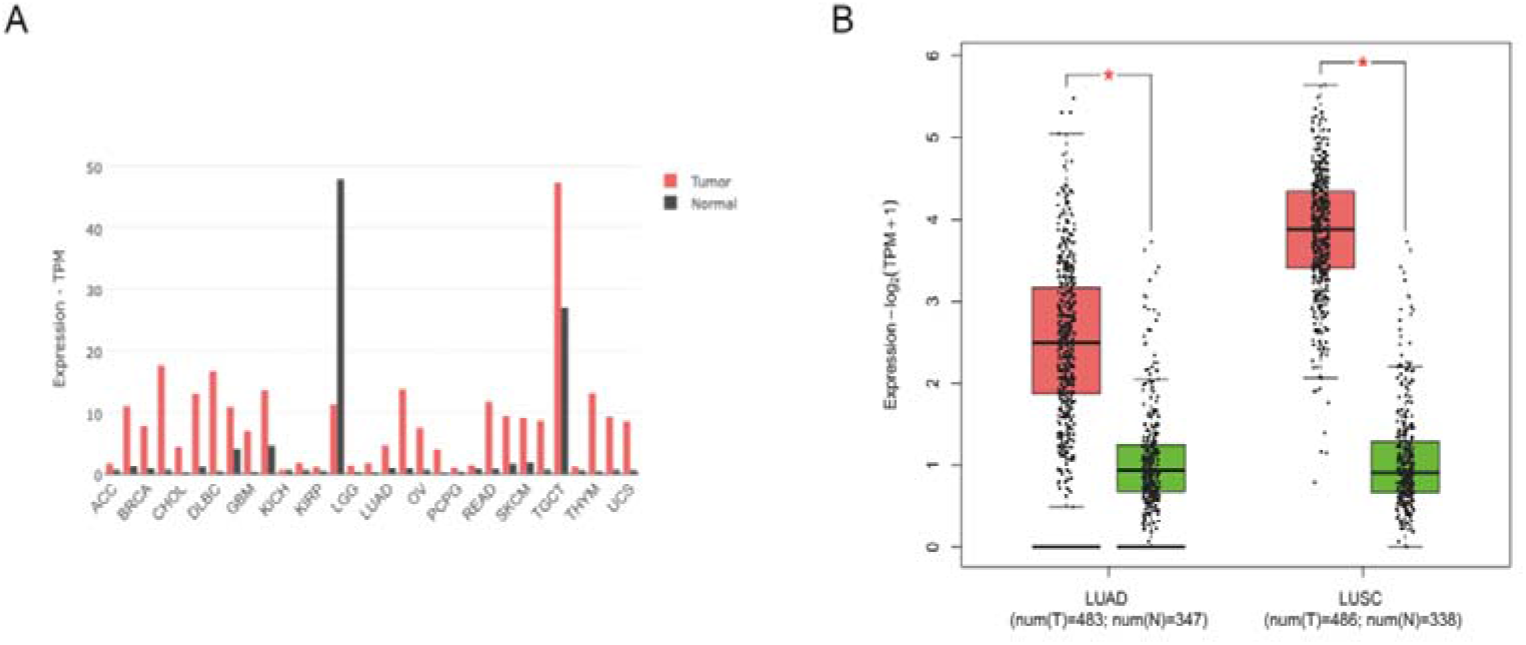
Expression of RAD51 mRNA in various cancers A:Expression of RAD51 mRNA in various cancers; B:Expression of RAD51 mRNA in lung adenocarcinoma and squamous cell carcinoma. (*p<0.05)

### 2.2 Relationship between RAD51 and prognosis of NSCLC patients

In lung adenocarcinoma, overall survival time was significantly reduced in patients with high RAD51 expression compared to those with low RAD51 expression (p=0.0026). In lung squamous cell carcinoma, RAD51 expression level had no significant effect on overall survival time (p=0.76) (Figure 2).

**Fig. 2.**
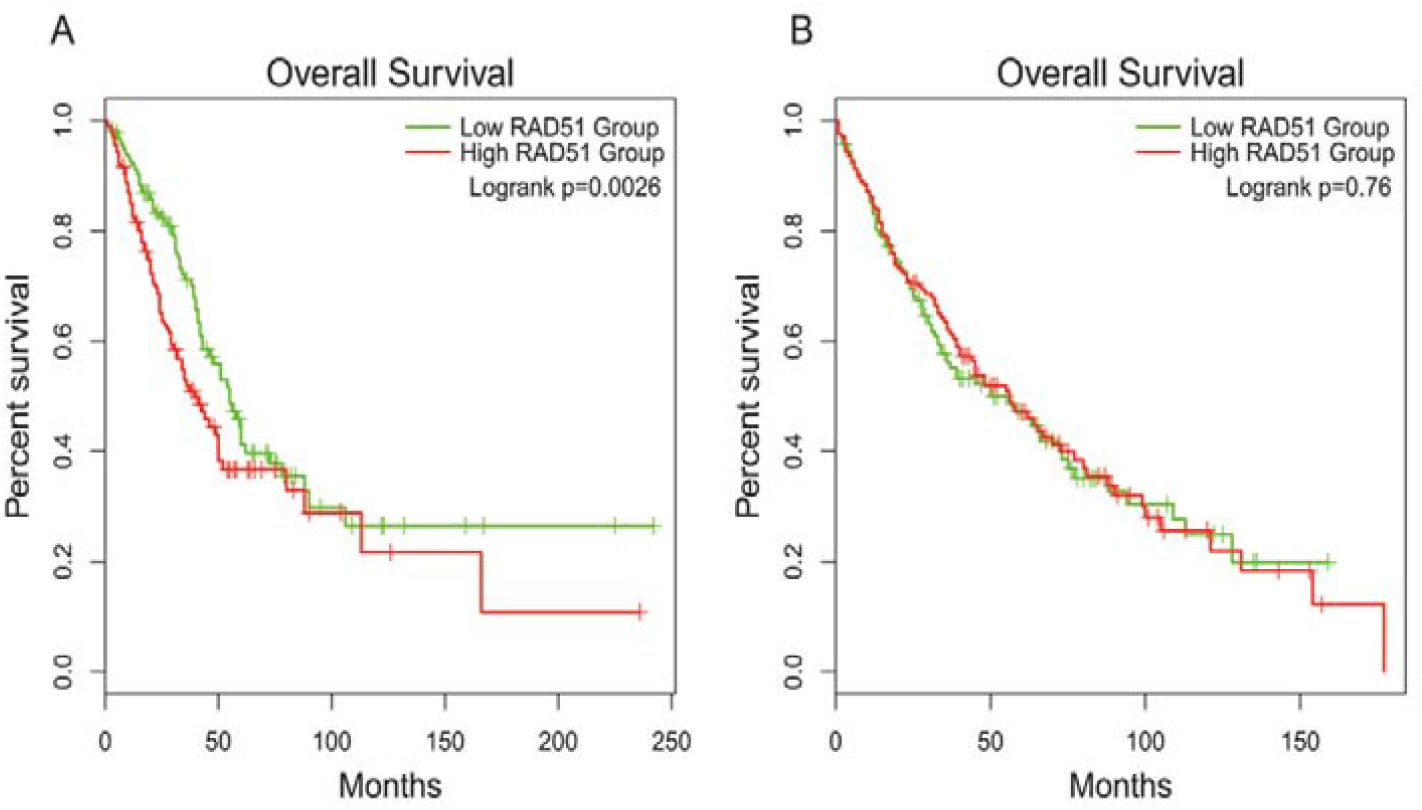
The relationship between RAD51 and prognosis of patients with non-small cell lung cancer A:The relationship between RAD51 and prognosis of patients with lung adenocarcinoma;B: The relationship between RAD51 and prognosis of patients with lung squamous cell carcinoma

### 2.3 Relationship between RAD51 and NSCLC stage

According to the latest international TNM staging standard for lung cancer (8th edition), lung cancer was divided into stage I (T1a, T1b, T1c), stage II (T2a, T2b), stage III and stage IV. The microarray analysis of RAD51 mRNA expression level in NSCLC patients with different pathological stages was conducted using GEPIA database, and the analysis results indicated that there were statistically significant differences in the expression level of RAD51 mRNA in lung adenocarcinoma patients with different pathological stages (p< 0.001), there was no significant difference in RAD51 mRNA expression level between lung squamous cell carcinoma and different pathological stages (p> 0.05). (In lung adenocarcinoma: F-value obtained by One-Way-ANOVA analysis was 5.98, and its corresponding P-value was 0.000528; In lung squamous cell carcinoma, the F value obtained by One-Way-ANOVA analysis was 1.16, and the corresponding p value was 0.326) (Figure 3).

**Fig. 3.**
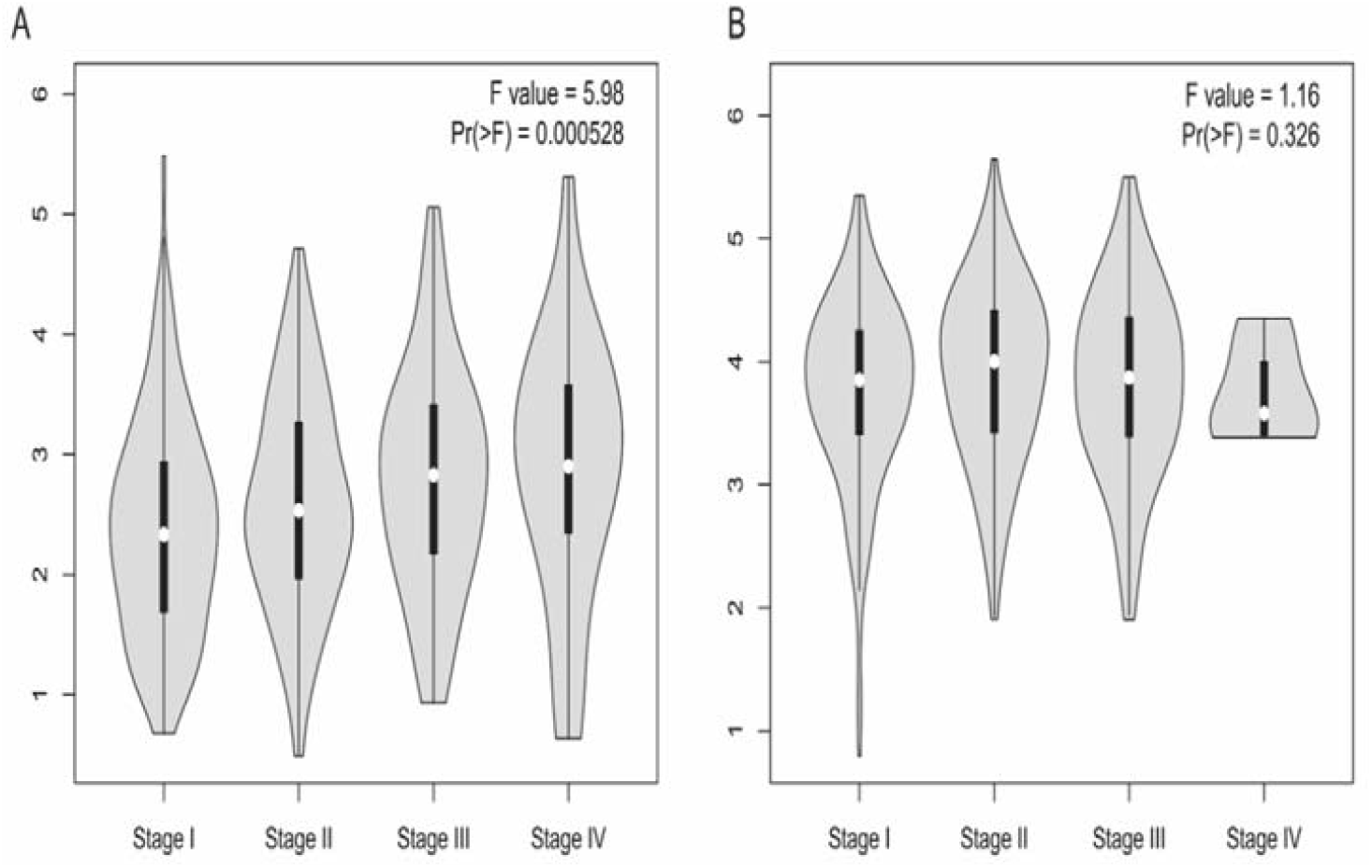
Relationship between RAD51 and staging of non-small cell lung cancer A:Relationship between RAD51 and staging of lung adenocarcinoma; B:Relationship between RAD51 and staging of lung squamous cell carcinoma

### 2.4 Correlation analysis of RAD51 with TP53, KRAS, EGFR, BRAF, MYC and ALK

The correlation between RAD51 and TP53, BRAF, EGFR, MYC, PD-L1 and KRAS was analyzed by GEPIA database. The analysis results indicated that RAD51 was positively correlated with TP53, BRAF, EGFR, MYC, PD-L1 and KRAS in NSCLC, and the correlation between RAD51 and MYC was the most significant, with an R value of 0.44. The difference was statistically significant (p< 0.001) (Figure 4).

**Fig. 4.**
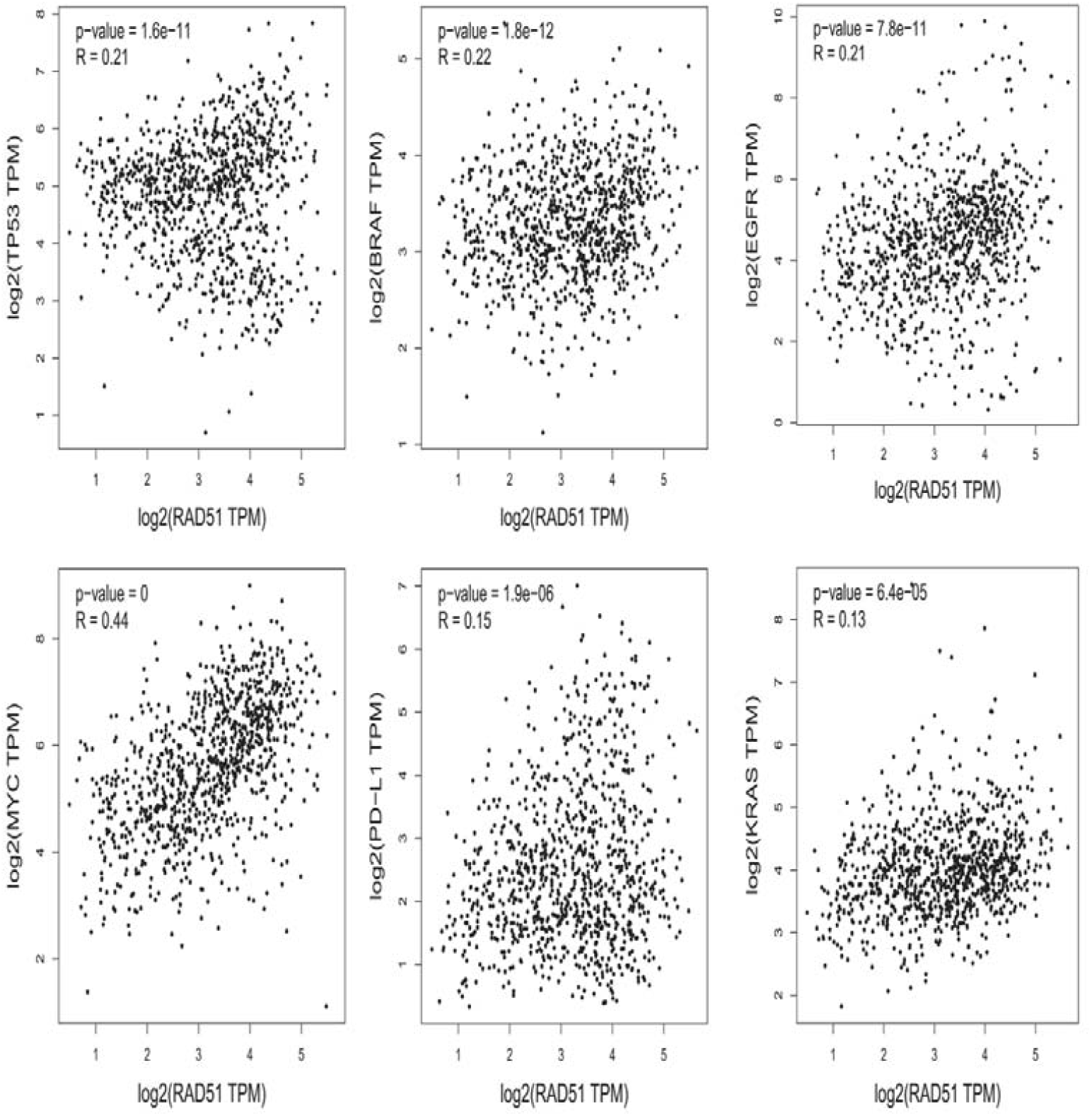
Correlation between genes and RAD51 in non-small cell lung cancer

### 2.5 Correlation analysis of RAD51 in NSCLC and immune cells in tumor microenvironment

For data analysis, purity of tumors was the main confounding factor in this analysis, and purity was corrected through spearman analysis. In lung adenocarcinoma, lung cancer cells were positively correlated with CD4+ memory T cells, CD8+T cells and M1 macrophages (p< 0.001) and were negatively correlated with regulatory T cells (p< 0.05), there was no correlation with M2-type macrophages (p> 0.05), the difference was statistically significant. In lung squamous cell carcinoma, tumor cells were positively correlated with M1-type macrophages (p< 0.05), had no correlation with CD4+T memory cells, CD8+T cells, M2 macrophages and Tregs cells (p> 0.05) (Figure 5).

**Fig. 5.**
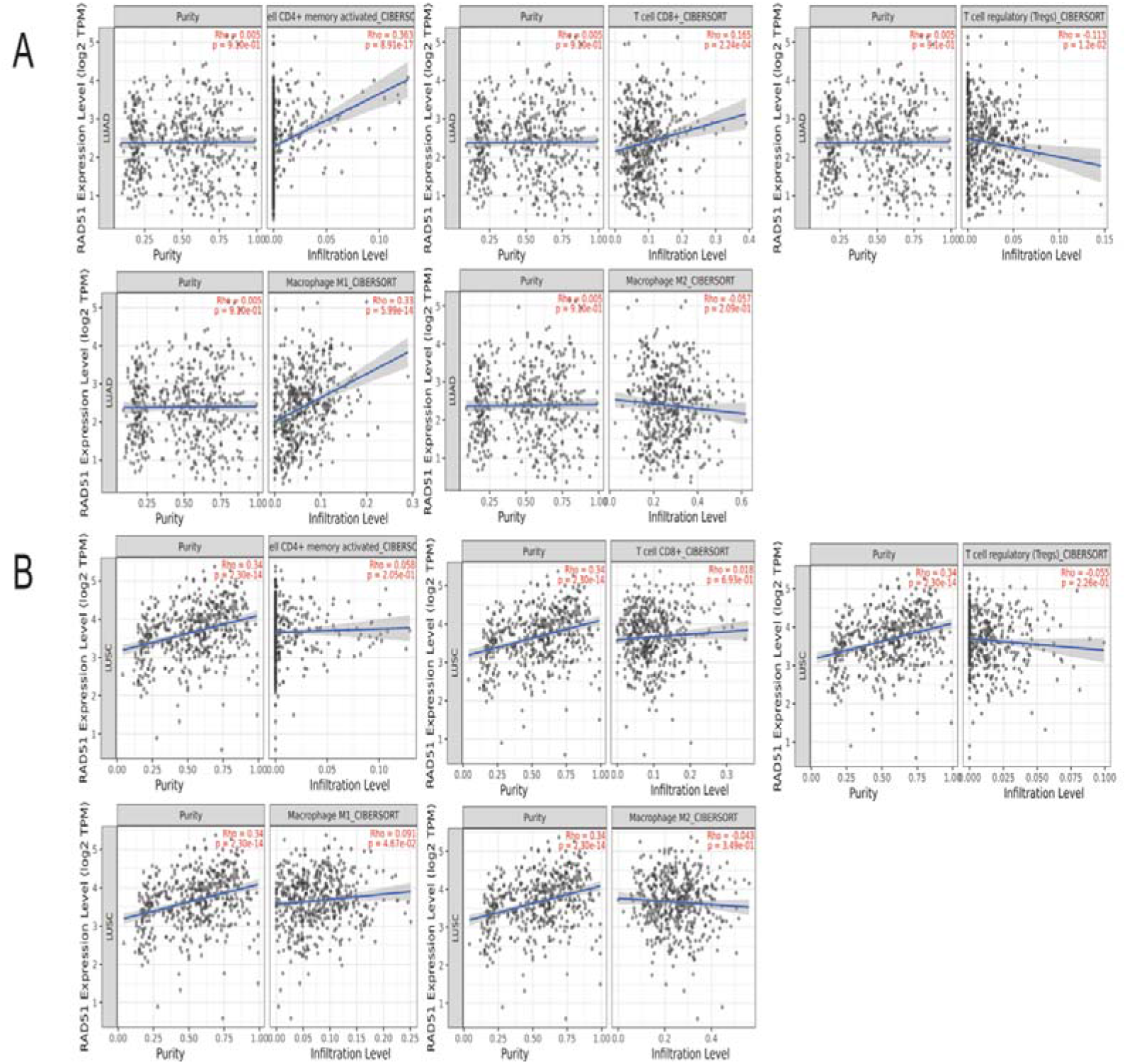
Relationship between RAD51 and immune cells in tumor microenvironment of non-small cell lung cancer A:The relationship between RAD51 and immune cells in the tumor microenvironment of lung adenocarcinoma; B:The relationship between RAD51 and immune cells in the microenvironment of lung adenosquamous tumor.

### 2.6 Protein expression difference of RAD51 in different normal tissues and NSCLC

The protein expression difference of RAD51 gene in human normal lung tissue and NSCLC was obtained from the HPA database, and the immunohistochemical staining results indicated that RAD51 was low expressed in normal lung tissue and high expressed in lung adenocarcinoma and lung squamous cell carcinoma (Figure 6).

**Fig. 6.**
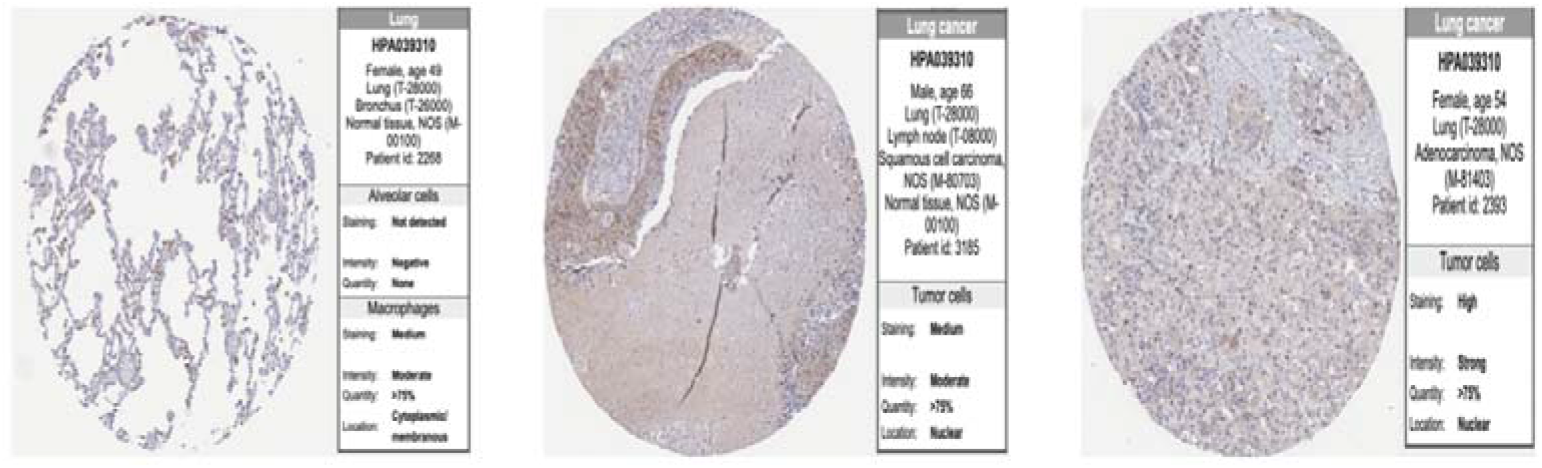
Expression of RAD51 in normal lung tissue and lung tissue of non-small cell lung cancer

## 3 Discussion

Worldwide, lung cancer is the most common cancer, the leading cause of cancer death in men, and the second leading cause of cancer death in women. Research by the World Health Organization (WHO) has shown that over the past few decades, The incidence of lung adenocarcinoma in males and females increases more rapidly than squamous cell carcinoma [14]. Smoking is the most commonly known cause of lung cancer, followed by environmental pollution, occupational exposure, and genetic factors [2, 4]. With the use of tyrosine kinase inhibitors in patients with EGFR, ALK, ROS1 and NTRK mutations, the treatment of lung cancer has made great progress. At the same time, the use of immune checkpoint inhibitors (ICIs) has greatly promoted the therapeutic prospects of NSCLC. ICIs are part of the current treatment of first-line NSCLC. For patients with stage III unresectable NSCLC, monotherapy, combined chemotherapy, or radiotherapy followed by definite chemoradiotherapy have all achieved good therapeutic effects [15]. The expression of programmed cell death protein-ligand 1 (PD-L1) in malignant tumor cells has become a potential biomarker for ICIs therapy [16]. In addition to PD-L1, the identification of other accurate and predictive biomarkers is still essential for the selection of appropriate ICI treatment. The preferred treatment for NSCLC is radiotherapy and/or cisplatin chemotherapy; however, a large number of patients diagnosed with NSCLC are at risk of organ metastasis and resistance to radiotherapy/chemotherapy, leading to treatment failure [17]. Therefore, it is necessary to further study the causes of lung cancer treatment failure and provide a more accurate basis for lung cancer treatment.

RAD51 is located on chromosome 15q15.1 and contains 339 amino acids. The sequence homology of vertebrate RAD51 with yeast and plants is 74%, and that of mouse RAD51 is 99%[18]. RAD51 can form nucleoprotein filaments on single-stranded DNA and has the function of discovering and entering homologous DNA sequences to realize accurate and timely DNA repair. DNA damage is induced by various endogenous and exogenous factors such as reactive oxygen species produced by cell metabolism, ultraviolet light, and genetically toxic chemicals. After DNA damage occurs, the repair mechanism is activated, which differs from double-strand DNA damage to homologous recombination. HR), non-homologous end joining (NHEJ), and microhomology-mediated end joining (MMEJ) [19]. HR involved by RAD51 is the most important mechanism of DNA double-strand repair and also one of the reasons for inducing tumor tolerance to radiotherapy and chemotherapy [20]. RAD51 is involved in the repair of DNA breaks induced by chemical inducers and mediates etoposide tolerance in lung cancer through HR and DNA-PKCs-dependent NHEJ [7]. RAD51 induces chemotherapy tolerance in breast cancer and chronic lymphocytic leukemia by activating HR [8, 9]. RAD51 promotes aerobic glycolysis to promote the development of pancreatic cancer, and inhibition of RAD51 can enhance the blocking of cellular glycolysis and improve the therapeutic effect on leukemia [10, 11]. More and more evidence shows that RAD51 is involved in tumor metabolism. The role and mechanism of RAD51 in NSCLC metabolism need further elaboration. In addition, RAD51 has been shown to promote epithelial-mesenchymal transition in esophageal squamous cell carcinoma and prostate cancer via p38/Akt/Snail and EGFR-Erk1/2/Akt signaling pathways. EMT) and metastasis [12, 13]. RAD51 mediates tumor metastasis through various mechanisms and will become a new therapeutic target to inhibit the progression of NSCLC and increase sensitivity to radiotherapy/chemotherapy.

We mined the expression of RAD51 in NSCLC based on the TCGA database and found that RAD51 was highly expressed in a variety of tumors (Figure 1A). In both lung adenocarcinoma and lung squamous cell carcinoma, the expression of RAD51 was higher than that in the normal control group (FIG. 1B). Analysis of age, sex and race showed that the occurrence of lung adenocarcinoma was independent of age, sex, and race (p> 0.05) (Table 1), the incidence of lung squamous cell carcinoma was associated with males (p< 0.05) (Table 2), possibly because there are significantly more males than females in lung cancer patients, and the proportion of males smoking is also significantly higher than females [21]. Survival analysis of patients with high and low expression of RAD51 showed that high expression of RAD51 predicted significantly shortened survival in lung adenocarcinoma (p< 0.01) (FIG. 2A), but had no significant effect on the survival cycle of patients with lung squamous cell carcinoma (p> 0.05) (Figure 2 B). Consistent with the research results of Li-Wei Zhang et al. [22], RAD51 expression was increased in lung adenocarcinoma, and the ubiquitination degradation of RAD51 was promoted by tripartite motif family proteins36 (TRIM36). Increase the radiotherapy sensitivity of lung adenocarcinoma. Our study found that the expression level of RAD51 mRNA in NSCLC was statistically significant in patients with lung adenocarcinoma at different pathological stages (p< 0.001) (FIG. 3A), there was no statistically significant difference in lung squamous cell carcinoma (p> 0.05) (FIG. 3B), consistent with the results of survival analysis of patients, suggesting that RAD51 has a significant carcinogenic effect on lung adenocarcinoma. We evaluated the difference in the expression of RAD51 protein in normal lung tissue and lung cancer using the HPA database and found that RAD51 was highly expressed in both lung adenocarcinoma and lung squamous cell carcinoma, and was strongly positive in lung adenocarcinoma (Figure 6).

tumor microenvironment (TME) is composed of various cells, extracellular components, and vascular networks. TME not only plays a key role in the occurrence, development, and metastasis of tumors but also has a profound influence on the therapeutic effect. Tme-mediated drug resistance is the result of crosstalk between tumor cells and their surrounding stroma [23]. Immune cells in TME include innate immune cells and adaptive immune cells, which can interact directly with tumor cells or enter into TME to perform biological functions through signal transduction such as chemokines and cytokines. Immune cells in Tmes can support or obstruct therapy and can change their activation status within Times [24]. tumor-associated macrophages (TAMs) are the key regulatory factor in the treatment of TME, which is associated with cancer prognosis [25]. Macrophages can be divided into M1 and M2 subtypes according to their polarization status [26]. M1 macrophages are believed to have antitumor effects and can recruit cytotoxic T lymphocytes (CTLS) to activate adaptive immune responses. M2 macrophages promote tumorigenesis and induce immune tolerance, and M1 and M2 have plasticity and mutual transformation [27, 28]. In NSCLC, extracellular vesicles secreted by tumor cells miR-21-5p, TRIM59, PKM2 and lncRNA TUC339 reshape the tumor microenvironment, promote tumor growth, proliferation, angiogenesis and M2 polarization, and mediate lung cancer liver metastasis [29-34]. CD4+T cells can target and kill tumor cells by releasing perforin, granase and other substances. In this study, CIBERSORT combined with TIMER database analysis showed that RAD51 was positively correlated with CD4+T cells, CD8+T cells and M1 macrophages in lung adenocarcinoma (Figure 5A). The reason may be that the elevated expression of RAD51 in lung adenocarcinoma stimulates the proliferation and differentiation of the above cells and reshaps TME. However, the relationship between T cells and RAD51 in tumor has not been reported. Therefore, it is necessary to further study the mechanism of RAD51 on TME, so as to provide more scientific basis for clinical tumor immunotherapy.

TP53, BRAF, EGFR, MYC, PD-L1 and KRAS have all been confirmed to be involved in tumor proliferation, migration, invasion and metastasis, and their mutation or high expression indicates poor prognosis of patients [35-37]. According to GEPIA database analysis, RAD51 was positively correlated with TP53, BRAF, EGFR, MYC, PD-L1 and KRAS in NSCLC, and the correlation between RAD51 and MYC was the most significant (p< 0.001) (Figure 4). Compared with wild-type lung cancer cells, RAD51 expression is higher in KRAS mutation, and MYC plays a key role in RAD51 overexpression. Inhibition of RAD51 can lead to enhanced DNA double-strand breaks, colony formation defects and cell death. It is suggested that RAD51 may be an effective therapeutic target to overcome chemical/radioresistance in KRAS mutated cancers [38].

In summary, in this study, the expression level of RAD51 mRNA in NSCLC tissues was mined through the TCGA database, and it was found that the high expression level of RAD51 in NSCLC was correlated with the overall survival of lung cancer. The survival cycle of lung adenocarcinoma patients with high expression of RAD51 was significantly shortened. Further analysis of lung cancer pathological tissue showed that the expression of RAD51 protein was positive in lung cancer. This study provides a theoretical basis for further exploring the pathogenesis, diagnostic markers and therapeutic targets of lung cancer. The limitation of this study lies in that it simply used TCGA and other databases for analysis, and did not provide further in vitro and in vivo experiments to further clarify the specific mechanism of action of RAD51 in lung cancer.

## Ethical Statement

This study does not contain any studies with human participants or animals performed by any of the authors.

